# Rapid 3D-STORM imaging of diverse molecular targets in tissue

**DOI:** 10.1101/2021.08.25.457670

**Authors:** Nicholas E. Albrecht, Danye Jiang, Robert Hobson, Colenso M. Speer, Melanie A. Samuel

**Author notes:** **Competing Financial Interests.** N.E.A, D.J., C.M.S., and M.A.S declare no competing financial interests. R.H. is an employee of Bruker Nano Surfaces.

## Abstract

The precise organization of fine scale molecular architecture is critical for the nervous system and other biological functions and would benefit from nanoscopic imaging methods with improved accessibility, throughput, and native tissue compatibility. Here, we report RAIN-STORM, a rapid and scalable imaging approach that enables three-dimensional nanoscale target visualization for multiple subcellular and intracellular targets within tissue at depth. RAIN-STORM utilizes conventional tissue samples, readily available reagents in optimized formulas, requires no specialized sample handling, and is suitable for commercial instrumentation. To illustrate RAIN-STORM’s ability for quantitative high-resolution nanoscopic tissue imaging, we utilized the well-organized but structurally complex retina. We show that RAIN-STORM is rapid and versatile, enabling 3D nanoscopic imaging of over 20 distinct targets to reveal known and novel nanoscale features of synapses, neurons, glia, and vascular. Further, imaging parameters are compatible with a wide range of tissue sources and molecular targets across a spectrum of biological structures. Finally, we show that this method can be applied to clinically derived samples and reveal the nanoscale distribution of molecular targets within human samples. RAIN-STORM thus enables rapid 3D imaging for a range of molecules, paving the way for high throughput studies of nanoscopic molecular features in intact tissue from diverse sources.

## Introduction

The advent of single-molecule localization microscopy (SMLM) techniques have greatly increased the ability to resolve the location, density, and nanoscale spatial relationships of diverse molecules (Bates et al., 2007). Yet, largescale adoption of these techniques for tissue analysis remains limited, in part because most SMLM approaches are challenging to apply to thick samples where aberrations and background can limit imaging. As a result, most analysis of tissue molecular architecture continues to rely on immunofluorescence microscopy, immunoelectron microscopy, and fluorescent protein reporters. These powerful tools have shown that cellular and tissue function intimately depends on small-scale arrangements of proteins and cells, but how nanoscale biological structures are organized, and their relative molecular composition, remain unknown for most targets, cell types, and tissue sources.

To help address this challenge a number of bespoke SMLM solutions have been developed. These include in-house, specialized microcopy and optical systems, customized analysis software, and advanced sample preparation techniques. For example, cultured cells have been visualized in three-dimensional volumes using astigmatism (Huang et al., 2008), biplane imaging (Juette et al., 2008), an engineered point spread function (Pavani et al., 2009), and 4pi-imaging (Bewersdorf et al., 2006), while tissue has been visualized using adaptive optics (Mlodzianoski et al., 2018), biplane imaging (Bewersdorf et al., 2006), self-interfering point spread functions (Bon et al., 2018), light-sheet approaches (Greiss et al., 2016), and ultrathin physical sample sectioning (Sigal et al., 2015). While these methods have greatly advanced 3D SMLM capability in the field and led to numerous discoveries (Bowler et al., 2019; Chamma et al., 2016; Leterrier et al., 2015; Sigal et al., 2015; Suleiman et al., 2013; van den Dries et al., 2013), significant challenges remain. The first is accessibility. Most 3D SMLM approaches rely on techniques, systems, and expertise that are available only to a handful of specialists. The second is simplicity and throughput. For instance, one useful approach called serial-section STORM uses ultrathin sectioning and reconstruction, but the labor and imaging time for this technique make it best suited for deep imaging of small sample numbers. The third is target compatibility within native tissue environments. Most current methods report imaging capabilities using only a small number of antibodies (Mikhaylova et al., 2015; Mönkemöller et al., 2015), and approaches such as expansion microscopy are not always suitable for low density targets (Chen et al., 2015; Ku et al., 2016; Tillberg et al., 2016).

We set out to develop a new imaging method that would complement and combine the strengths of currently available methods for nanoscopic tissue imaging in general and for neural circuits specifically. Our criteria for this method were to improve the accessibility, throughput, and compatibility of 3D nanoscopic tissue imaging so that it can be broadly and readily applied to diverse tissue sources and target types. Toward this goal, we present <u>Ra</u>pid Imaging of tissues at the <u>N</u>anoscale (RAIN-STORM), a method for utilizing standard tissue samples to generate SMLM data at depth for a wide range of molecular targets using commercially available reagents and imaging systems. To achieve this goal, we optimized labeling and imaging conditions from 125 distinct tested parameters for over 20 molecular targets in murine tissue and validated methods in four additional species. We show that RAIN-STORM is rapid and versatile, enabling 3D nanoscopic imaging with a 24-hour turnaround. Further, these imaging parameters are compatible with a wide range of tissue sources and a large number of molecular targets across a spectrum of biological structures. Finally, we show that this method can be applied to clinically derived samples and reveal the nanoscale distribution of molecular targets within human samples. RAIN-STORM thus enables rapid 3D STORM for a range of molecules, paving the way for high throughput studies of nanoscopic molecular features in intact tissue from diverse sources.

## Results

A central criterion for our method is compatibility with standard tissue preparation protocols that require little special tissue handling. To test this, we used the mouse retina because it has a highly defined laminar structure that provides endogenous fiduciaries for evaluating the labeling precision of multiple molecular targets (Sanes & Zipursky, 2010). To begin, retinas were harvested from wildtype adult mice and processed with a standard tissue preparation protocol (Albrecht et al., 2018) that involves a short primary fixation, cryoprotection, embedding in tissue freezing medium, sectioning at 10 µm on a cryostat, and mounting on coated glass slides (**Figure 1a**). Using this basic framework, we tested and adapted variations for each preparatory step to find ideal conditions suitable for SMLM-based imaging. Sample preparation thus does not require labor-intensive handling, and results in staining ready slides within hours of tissue harvest. For turnkey nanoscopic imaging, we utilized a commercial SMLM imaging system, the Vutara 352 (Bruker, Billerica MA), together with associated commercial image analysis software. This system enables multicolor STORM imaging within a 40µm-by-40µm planar region of interest and achieves single molecule imaging of individual emitters by recording the point spread function (PSF) in two imaging planes simultaneously (**Figure 1 – figure supplement 1**). Single-molecule localization in 3D is based on calibration data generated from fiducial imaging.

**Figure 1:**
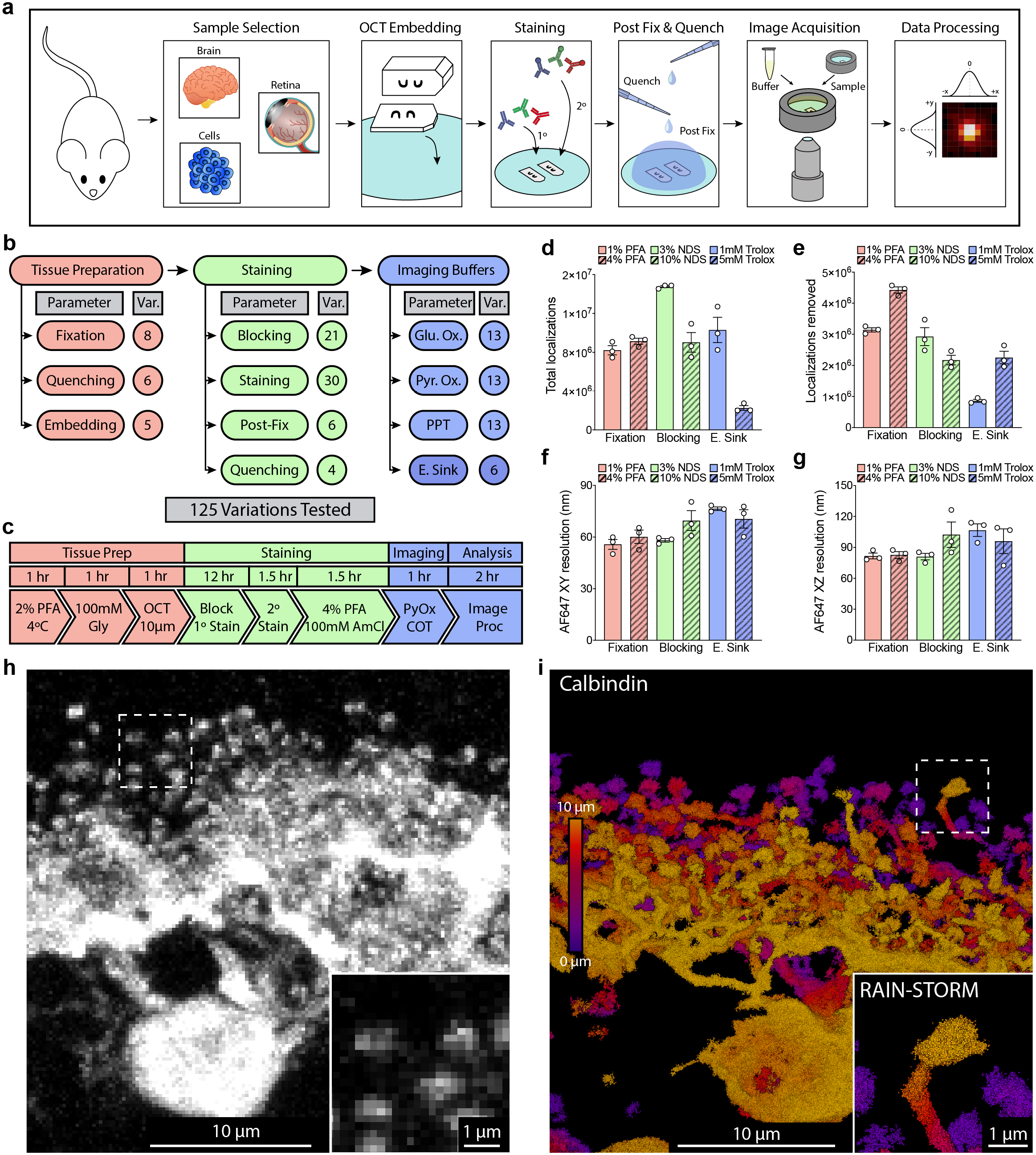
Nanoscale 3D imaging of neurons in tissue. **a**, Schematic of the RAIN-STORM workflow. Eyes are enucleated, and retina cups are dissected, fixed, and quenched. Retinal cups are then cryopreserved and embedded in OCT prior to sectioning and mounting onto prepared coverslips. Samples are stained with primary and secondary antibodies, post-fixed and quenched, and finally mounted in open wells containing imaging buffer. **b**, Schematic of tested parameters with the number of variations tested for each category. **c**, Optimized RAIN-STORM parameters and associated preparation timeline. **d-g**, Representative quantifications for six exemplar tested parameters representing three different stages of sample preparation, staining, and imaging. The total localizations acquired (**d**) impacts the final localization density, while the total removal of non-structured localizations (**e**) measures the noise/background for each condition. Calculated resolutions are also provided for the XY-plane (**f**) and XZ-plane (**g**) for each condition using FRC. Among these conditions, low levels of PFA and Trolox and moderate levels of normal donkey serum provided the best image quality metrics and the least unwanted signal. **h**, Unoptimized STORM image of a horizontal cell labeled with Calbindin. Putative synapses are indistinct with little defining morphology or contiguous structure. **i**, RAIN-STORM image of a Calbindin-labeled horizontal cell using optimized parameters demonstrates clear structural detail across the neuronal arbor. Distinct synaptic terminals are visible together with the connecting stalk arising from the neuron. N = 3 animals. Scale bars = 10 and 1 µm. Data are represented as the mean ± the s.e.m.

We selected the calcium buffering protein Calbindin (Calb1) and the synapse protein PSD95 (Postsynaptic density protein 95, also known as Dlg4) to optimize our labeling and imaging parameters. Since Calbindin specifically and densely fills the cell body and neurite terminals of retina horizontal neurons (Celio, 1990; Uesugi et al., 1992), while PSD95 moderately labels synapses (Hunt et al., 1996; Koulen et al., 1998), we reasoned that optimizing our imaging processes around these targets would enable us to evaluate the efficacy of our method across a range of cell structures and protein densities. Moreover, because Calbindin and PSD95 are found in other central nervous system regions (Celio, 1990; Hunt et al., 1996) preparation methods compatible with these targets may extend to other tissue types.

Significant image aberration and background in STORM tissue imaging can result from suboptimal sample preparation, imaging buffer, and image acquisition conditions. We reasoned that tuning these parameters for 3D STORM imaging could markedly improve nanoscale image quality. We focused on the following features: preservation of tissue structure, epitope integrity, reduced background fluorescence, high labeling specificity, uniform antibody penetration, and increased localization density within samples. We performed STORM imaging following precise manipulation of 80 fixation and staining variations and 45 buffer conditions, resulting in a total of 125 test conditions (**Figure 1b**, **Supplemental Table 1**, and **Figure 1 – figure supplement 2**). We chose the best performing conditions based on the following criteria: 1) The total number of localizations acquired (**Figure 1d**), 2) epitope and morphological detail preservation, 3) the relative background level (noise) observed as isolated localizations or spurious antibody signal (**Figure 1e**), and 4) the calculated resolution across planar (XY, **Figure 1f**) and axial (XZ, **Figure 1g**) image dimensions using Fourier Ring Correlation (FRC) metrics (Nieuwenhuizen et al., 2013). Samples were obtained from three independently prepared animals, giving three independent measurements per condition tested. Within this parameter space, we first investigated the effects of primary fixation on tissue quality and imaging. We found that autofluorescence could generally be lowered by reducing fixative concentrations, and that temperature played an important role in limiting structural distortions and preserving epitopes. A cold fixation (4°C) at a relatively low concentration of paraformaldehyde (2% PFA) yielded the best labeling density and imaging metrics (**Supplemental Table 1** and **Figure 1 – figure supplement 2**). Second, we tested diverse fixation quenching reagents, staining buffer components, primary and secondary antibody concentrations, and both post-fixation and post-fixation quenching methods. Of these, we found that variations in post-fixation and fixation quenching conditions had the largest effects on tissue integrity and final image quality (e.g., 47.6 ± 2.8 nm for those with post-fixation versus 72.8 ± 3.0 nm for those without, **Supplemental Table 1**). Image resolution was further increased, and background fluorescence decreased, by quenching in 100mM NH_4_Cl for 30min (50.5 ± 2.1 nm for treated versus 72.8 ± 3.0 nm for untreated samples, **Supplemental Table 1**)

We then tested multiple dilutions for a range of primary and secondary antibodies to identify the concentrations that resulted in the highest signal-to-noise ratio. (**Supplemental Table 1**). We found that many of our targets benefitted from using increased primary and secondary antibody concentrations relative to standard histological preparations (e.g., anti-Calbindin primary antibody is optimal at 1.0 ug/ml in STORM compared to 0.1 ug/ml in diffraction-limited microscopy). For secondary antibodies, we found a significant difference between conditions, with high concentrations (5.0 µg/ml) providing increased labeling density relative to standard confocal dilutions (0.5 µg/ml, ∼12.4×10^6^ vs. ∼7.2×10^6^, localizations respectively) though both were sufficient to image primary-labeled structures for all tested fluorophores (**Supplemental Table 1**).

Finally, we undertook a thorough examination of imaging buffer conditions and tested the effects of 1) oxygen-scavenging enzymes, 2) catalase concentration, 3) the ratio of thiols [ß-mercaptoethanol (BME) versus ethanolamine (MEA)], and 4) triplet-state quenching [cyclooctatetraene (COT)]. Results from these tests are summarized in **Supplemental Table 1**. For example, we found that the use of pyranose oxidase was preferable to glucose oxidase as it provided a more stable and longer-lived imaging environment. In addition, we determined that the addition of 2mM COT led to an increase in the total number of localizations across both channels (**Supplemental Table 1** and **Figure 1 – figure supplement 2**). The final sample preparation and imaging method was chosen from all tested parameters using the best performing parameters from each optimization step. This selection considered whether balanced data acquisition and quality could be achieved, the total number of localizations acquired, and the absolute image resolution for each fluorophore. Our final preparation and imaging method is termed RAIN-STORM and is comprised of readily accessible reagents and six simple steps that include a moderate initial fixation (2% PFA for one hour at 4°C), quenching of autofluorescence (100 mM Glycine for one hour at 4°C), primary and secondary staining using a serum-based blocking and permeabilization buffer (5% serum, 0.3% Triton-X100), post-fixation (4% PFA for 30 min at 4°C), and a final round of quenching (100 mM NH_4_Cl for 30min, **Figure 1b**).

To illustrate RAIN-STORM’s ability we reconstructed horizontal cells in 3D. We found our methods markedly improved quantitative resolution of nanoscopic cellular features (**Figure 1 h-i**). For example, we were able to observe distinct neurites of diverse morphologies arising from the cell body, which terminated in a high number of postsynaptic invaginations. These synaptic terminals were easily resolved and structurally separated from their neighbors, demonstrating markedly improved spatial resolution (31.2 ± 0.4nm and 61.1 ± 2.9nm for Calbindin labeled with CF568 and AF647, respectively, compared to ∼200-250nm resolution for typical confocal microscopy). RAIN-STORM also provides improved resolution at along the Z axis at depth, suggesting it is well suited for 3D super-resolution imaging of molecular targets within tissue volumes. To assess this, we computed the resolution of RAIN-STORM as a function of image depth at discrete imaging planes throughout the image volume. As expected, we found that planar resolution was improved closer to the objective relative to further away (30.0 ±1.5nm versus 42.3 ± 4.8nm**, Figure 1 – figure supplement 3**). To accommodate biological variability and nonhomogeneous structures, we report the aggregate resolution across the entire 10µm image stack. We achieve a Z-axis resolution of 42.2 ± 1.2nm and 79.3 ± 0.3nm for CF568 and AF647, respectively. RAIN-STORM thus offers the advantage of improved spatial resolution and can be used to visualize nanoscopic neural structures in large-field 3D volumes across the imaging plane to reconstruct all processes arising from a single cell (**Figure 1, Video 1**).

RAIN-STORM is also versatile, enabling nanoscopic imaging of a diverse array of molecular targets. To assess this, we took advantage of the wide spectrum of specific antibody markers available in the retina (Sanes & Zipursky, 2010) and tested the robustness of this approach across 22 validated molecular targets (**Supplemental Table 2, Figure 2** and **Figure 2 – figure supplement 1**). These included diverse cell structures, types, and molecular targets, such as synapse proteins (Bassoon, Bsn; RIBEYE, Ctbp2; Connexin 43, Gja1; Dystrophin, Dmd; Piccolo, Pclo; PSD95, Dlg4), vasculature markers (CD31, Pcam1; Collagen IV, Col4A1; Desmin, Des; CSPG4, Cspg4), glia (Iba1, Aif1; GFAP, Gfap; GS, Glul), excitatory interneurons (PKCα, Prkca; SCGN, Scgn), presynaptic photoreceptors (CAR, Arr3), and intracellular proteins and structures (Tau, Mapt; Tomm20, mitochondria, **Figure 2** and **Figure 2 – figure supplement 2**). We found that approximately 90% of the commercially available antibodies tested were compatible with RAIN-STORM and could be used to successfully obtain SMLM images of individual targets at depth within intact tissue slices (**Figure 2, Videos 1-3**). Further, in all cases labeling of the proper cell type or structure at the proper location was observed (**Figure 2 – Figure Supplement 1 and 2**). In addition, our results resolve novel nanoscale structural features and molecular distributions. For example, we observed that astrocyte end feet form fine (∼100-200nm) filamentous structures and what appears to be a mesh of fibers interdigitating the ganglion cell layer (**Video 3**). We also resolved individual dystrophin puncta in the outer retina synapse layer, which are thought to interact with actin filaments to enable contact formation between photoreceptors and ON-bipolar cells (Schmitz & Drenckhahn, 1997) (**Figure 2a**). Finally, we observed vascular associated interpericyte tunneling nanotubes (IP-TNTs, **Figure 2b**, **Video 2**). These ∼500nm diameter structures enable pericyte driven neurovascular coupling (Alarcon-Martinez et al., 2020) but their molecular composition is largely unknown. We show that Collagen IV comprises the core of these tubes, forming a solid filament connecting two hollow blood vessels (**Video 2**). These results suggest that RAIN-STORM is effective to image many targets, resolves biologically relevant protein localization, and can be used to discover unknown nanoscopic cellular structures with high fidelity.

**Figure 2:**
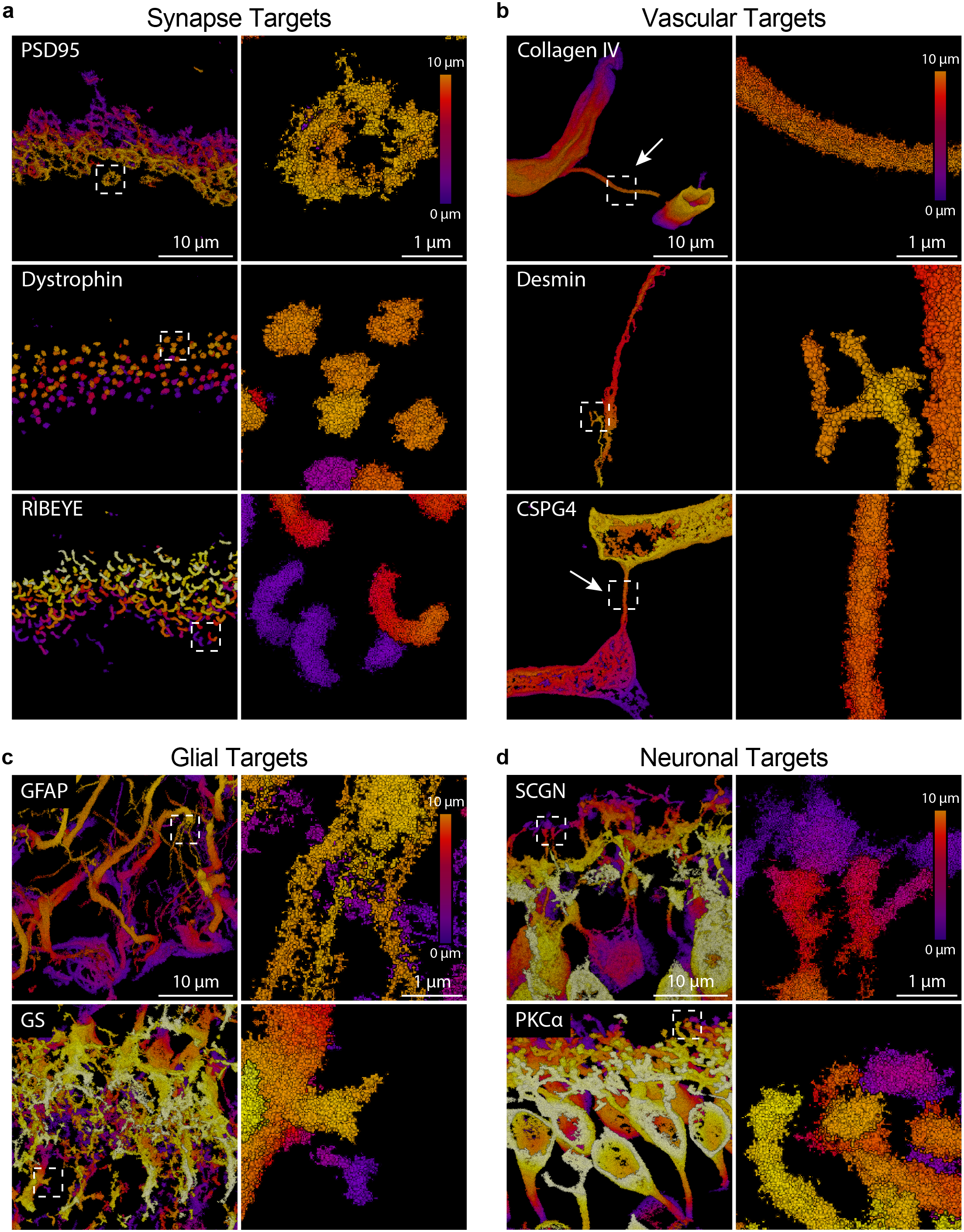
RAIN-STORM delivers robust imaging for a diverse array of molecular targets. **a**, RAIN-STORM imaging of synaptic proteins PSD95, a marker for photoreceptor terminals, dystrophin, a synaptic structural protein in the outer plexiform layer, and RIBEYE, a scaffolding protein present in ribbon synapses. Individual synapse terminals can be observed with all three markers. **b**, RAIN-STORM imaging of vasculature proteins Collagen IV, a marker for blood vessels, Desmin, a filament protein that marks subsets of pericytes and vascular associated smooth muscle cells, and CSPG4, a pericyte marker. Vasculature associated interpericyte tunneling nanotubes are visible with both Collagen IV and CSPG4 (arrows). **c**, RAIN-STORM imaging of glial proteins GFAP, a marker for astrocytes, and GS, a marker for Müller glia. In each case, fine features of these cell types can be observed, including filament-like protrusions from astrocytes. **d**, RAIN-STORM imaging of excitatory neurons, SCGN, a marker for subsets of cone bipolar cells, and PKCα, a marker for rod bipolar cells. In each case, fine features of these images are representative of those acquired from N = 3 animals. Scale bars = 10 and 1 µm.

Multicolor RAIN-STORM imaging is also robust across both multiple intracellular targets and in tissue from diverse species (**Figure 3**). We observed robust imaging with multiple target combinations. These include a neuron subtype marker together with synapse antibodies, staining for two distinct post-synaptic neuron types, as well as co-staining for vascular markers and astrocytes (**Figure 3a-c**). Using these combinations, we were able to resolve contact sites, overlapping and non-overlapping cellular structures, and fine cellular interactions. For example, astrocytes and blood vessel co-staining revealed fine astrocytic filaments enshrouding and forming close contacts with neighboring vessels (**Figure 3b**). Further, RAIN-STORM imaging was compatible with tissue from diverse species, revealing both conserved and unique nanoscopic features across mouse, rabbit, macaque, and pig tissues (**Figure 3d-i**). For instance, co-staining for rod bipolar cells and their synapses (PKCα and PSD95, respectively) in adult macaque, rabbit, and pig retina showed that despite similar functions and molecular identities, rod bipolar cells have incredibly diverse dendritic structures (**Figure 3d-f**). Rabbit rod bipolar cell bodies were rounder and less elongated relative to pig, macaque, and mouse, with arbors that branched closer to the cell body. Notably, PSD95 labeling was confined to the outer layer of rod bipolar dendrites across species, suggesting presynaptic contacts are restricted to this region independent of species type. These data indicate that RAIN-STORM is suitable for 3D multicolor imaging in tissue from a range of species to simultaneously visualize diverse molecular targets.

**Figure 3:**
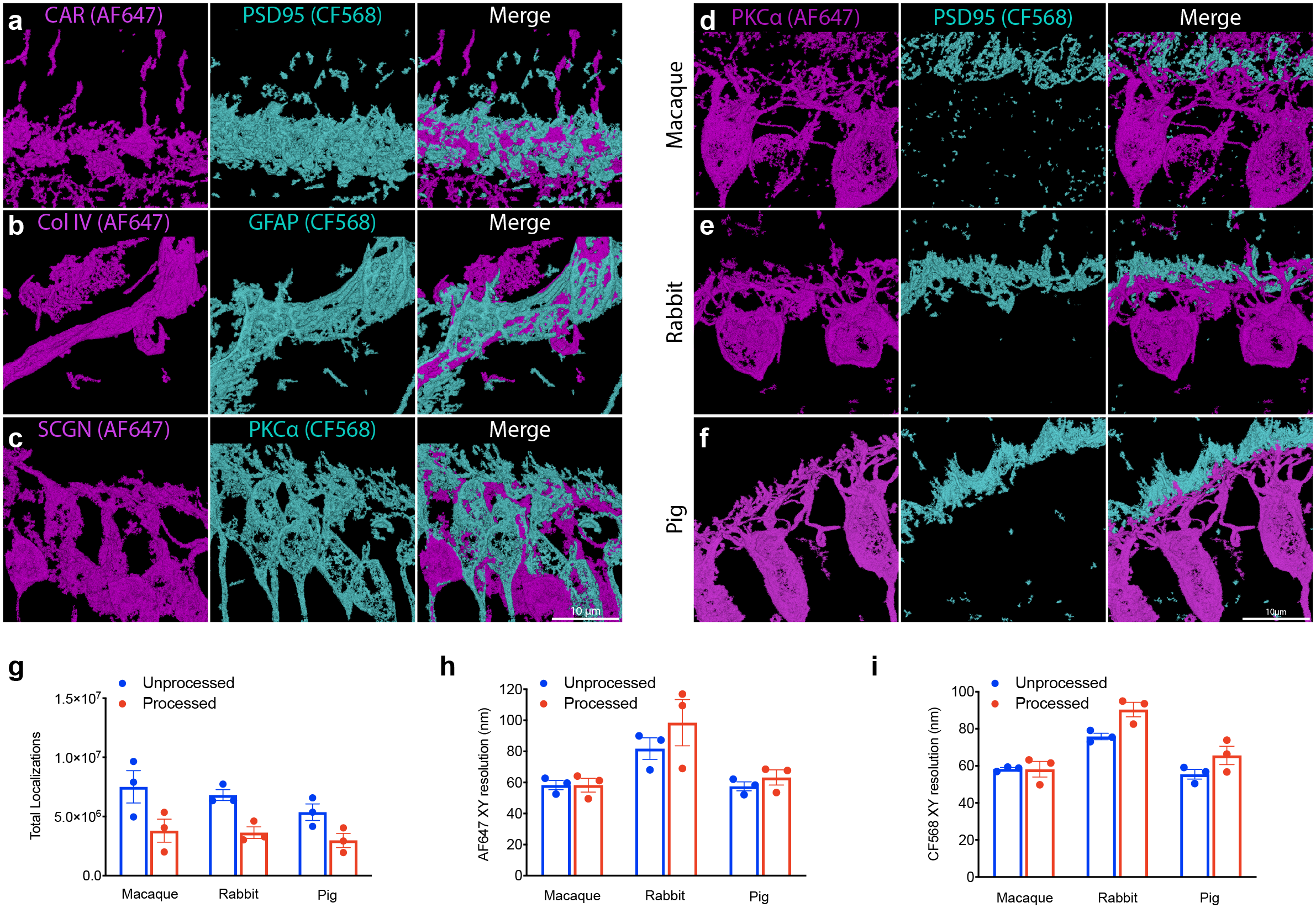
RAIN-STORM enables dual-channel super resolution imaging among diverse species. **a-c**, RAIN-STORM imaging of two independent molecular target, including **(a)** cones (CAR, magenta, AF647) and synaptic terminals (PSD95, cyan, CF568), **(b)** vasculature (Collagen IVa, magenta, AF647) and astrocyte (GFAP, cyan, CF568) interactions, and cone bipolar cells (SCGN, magenta, AF647) and rod bipolar cells (PKCα, cyan, CF568). **d-f**, RAIN-STORM imaging can be extended to diverse species. Retinas from macaque (N = 2 animals), **(e)** rabbit (N = 2 animals), and **(f)** pig (N = 3 animals) were labeled with antibodies to rod bipolar cells (PKCα, magenta, AF647) and rod photoreceptor terminals (PSD95, cyan, CF568). **g**, The total number of localizations that were acquired pre- and post-processing for images in **d-f**. **h** are displayed together with the XY planar resolutions of both AF647 (PKCα, magenta) and CF568 (PSD95, cyan). Scale bars = 10 and 1 µm. Data are represented as the mean ± the s.e.m.

RAIN-STORM can also be used to visualize, quantify, and measure features of small structures, including synapses. To assess this, we imaged retina ribbon synapses in the outer plexiform layer **(Figure 4a**). This region has two advantages. First, the architecture and composition of outer retina synapses have been well-characterized, allowing validation of our method. Second, because each photoreceptor forms connections at one distal location, the relationship between the structure and of both pre- and postsynaptic neurons relative to their connectivity can be directly examined. To resolve both murine ribbon synapses and their postsynaptic partners, we applied antibodies against the synapse scaffolding protein RIBEYE (Moser et al., 2020) together with the postsynaptic bipolar marker PKCα (**Figure 4a-c**) and used CF568- and AlexaFluor647 (AF647)-conjugated secondary antibodies to serially visualize both targets. We reconstructed individual RIBEYE labeled ribbons and assessed their shape, 2D-projected length, and 2D-projected area. RIBEYE showed a rich variety of ribbon morphologies, including many that appeared in a horseshoe shape and others that appear flatter, consistent with the more elongated contacts of basally located rods (Li et al., 2016) Measurements taken of individual ribbons (n = 844) across four adult mice showed an average ribbon length of 1.92 ± 0.30µm and a 2D projected area of 0.52 ± 0.13 µm^2^ **(Figure 4d-f** and **Video 4)**. Notably, these values are comparable to those obtained using Structured Illumination Microscopy (Dembla et al., 2020) (SIM, 1.25 ± 0.05 µm to 1.95 ± 0.07 µm in length), validating the specificity and accuracy of RAIN-STORM for measuring cellular structures. Together, these data suggest that RAIN-STORM can be used to reconstruct and measure defined neuron and synapse types at nanoscale resolution within tissue.

**Figure 4:**
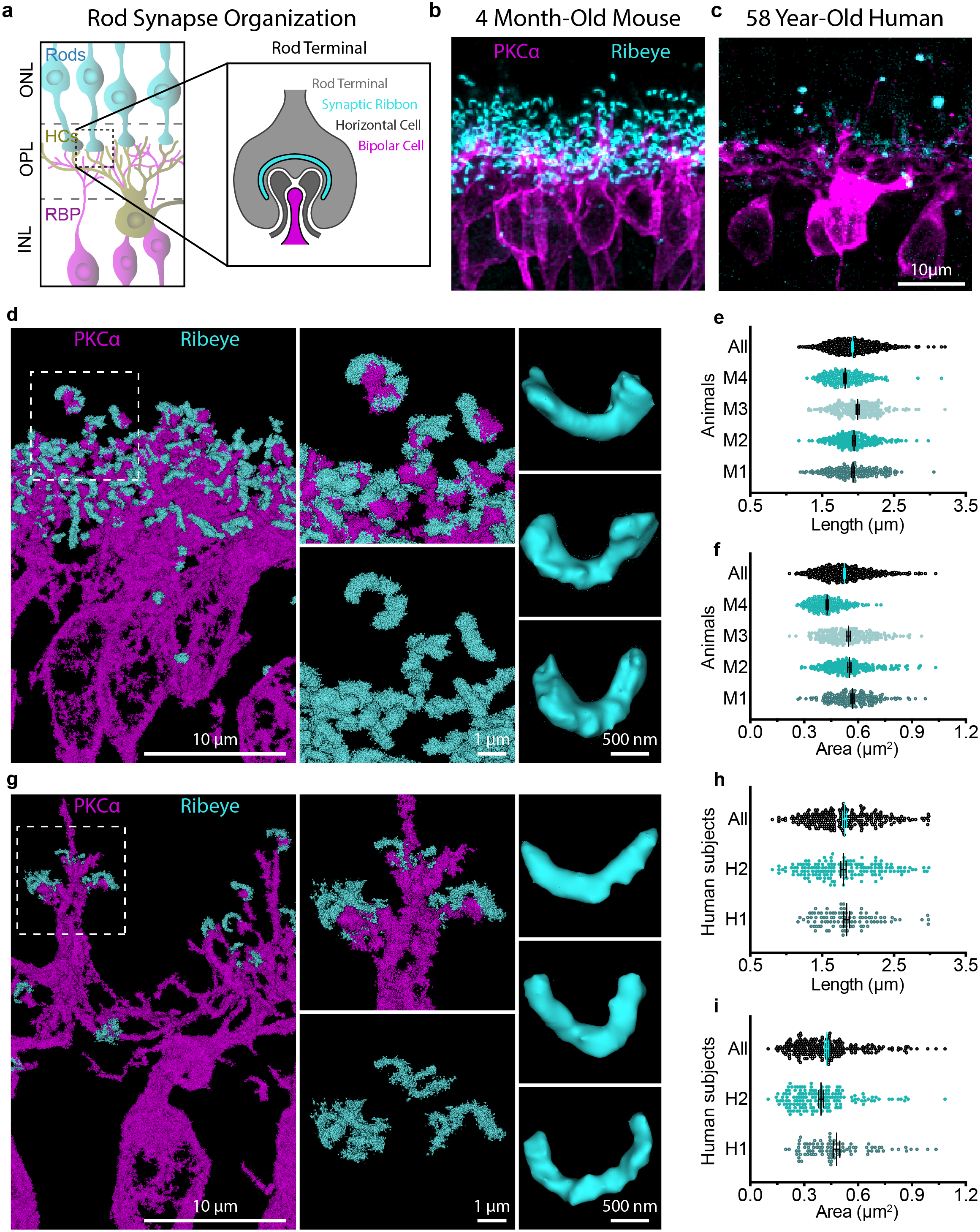
RAIN-STORM quantifies structural and molecular features of synapses. **a**, Schematic of retina outer plexiform synapse organization. Rod photoreceptors terminals (cyan) are presynaptic to invaginating post-synaptic horizontal cells (grey) and bipolar cells (magenta). **b-c**, Diffraction-limited imaging of murine (**b**) and human (**c**) retina outer plexiform synapses. Presynaptic photoreceptor terminals are labeled with RIBEYE (cyan) while postsynaptic bipolar cells are labeled with PKCα (magenta). **d**, Dual color RAIN-STORM imaging of murine rod bipolar cells (PKCα) and ribbon synapses (RIBEYE) allows individual quantification of ribbons and shows a rich variety of morphologies. **e-f**, Individual outer retina synapses (n = 844) were reconstructed from adult mice (N = 4), and the largest 2D projected length (**e**, 1.92 ± 0.30 µm) and total area (**f**, 0.52 ± 0.13 µm^2^) were quantified for each synapse. **g**, RAIN-STORM imaging of human rod bipolar cells (PKCα, magenta) and ribbon synapses (RIBEYE, cyan) resolves interactions between pre- and postsynaptic neurons. **h-i**, Individual outer retina synapses (n = 263) were reconstructed from human adult donors aged 40-58y (N = 2), and the largest 2D projected length (**h**, 1.81 ± 0.03 µm) and total area (**i,** 0.43 ± 0.01 µm^2^) were quantified for each synapse. Scale bars = 10 µm, 1 µm, and 500 nm. Data are represented as the mean ± the s.e.m.

Finally, we found that RAIN-STORM is compatible with human tissue. Ribbon synapses are also present in the human outer retina (Moser et al., 2020), but their relative size and organization have not been well mapped. Eyes from two adult human donors (aged 40 and 58 years) were processed for RAIN-STORM with similar parameters to those in mouse and stained with antibodies against RIBEYE and PKCα. As above, images were acquired using sequential imaging for each of the two channels. Reconstruction and quantification of human RIBEYE-labeled synapses showed an average length of 1.81 ± 0.03µm, which is similar in size to mouse ribbon synapses. Human ribbons also displayed a similar variety in shape and morphology as those found in mouse (**Figure 3g-i, Video 5**). Of note, we did observe fewer synapses in the human samples relative to the mouse, although this may be due to the inherent delay in human post-mortem sample collection rather than a biologically relevant difference.

## Discussion

In this paper we introduce RAIN-STORM, a rapid and scalable imaging approach that enables 3D nanoscale target visualization for multiple subcellular and intracellular targets within tissue at depth. We took advantage of the well-organized but structurally complex retina circuit to demonstrate that RAIN-STORM can resolve nanoscale features for a wide range of cell types and structures. This enabled us to validate known nanoscale structures as well as map novel cellular features of neurons, glia, and the vasculature. In addition, we visualized and quantified hundreds of single human and mouse synapses across multiple individuals. The acquisition of this dataset was facilitated by the high throughput nature of RAIN-STORM, and we show that this method is practical for analyzing specimens from multiple samples and species. Finally, because RAIN-STORM was developed to be compatible with a commercial imaging system and standard tissue processing, it is open to a range of researchers, applications, and clinical samples.

RAIN-STORM offers a number of advantages relative to existing 3D SMLM methods, including its experimental accessibility. We maximized its availability by using a user-friendly commercial imaging system so that this method could be readily adopted by the wider scientific community. With these advances, 3D tissue RAIN-STORM imaging has the capacity to be as routine as confocal imaging in many laboratories. In contrast, other 3D STORM imaging approaches, while excellent (Huang et al., 2008; Nehme et al., 2020; Punge et al., 2008; Xu et al., 2015), rely on custom built microscopes and require optics expertise and resources that are largely unavailable to most researchers. Also of note, the optimal R_X;Z_ resolution achieved with RAIN-STORM (R_X;Z_ = 61.1±2.9; 79.3±0.3) approaches, and in some cases exceeds, that of bespoke systems (e.g. SELFI, R_X;Z_ = 68±20; 115±32nm). RAIN-STORM is also rapid. We developed our method for use with conventionally prepared tissue samples, requiring only standard reagents and tools. Unlike serial array tomography EM (Micheva & Smith, 2007), serial section SMLM (Nanguneri et al., 2012) or related (Li et al., 2016) approaches, RAIN-STORM does not require time intensive ultrathin sectioning, successive section imaging, alignment, or reconstruction. Due to the straightforward nature of sample preparation and the relatively large field of view of our system, extensive datasets can be acquired quickly. For instance, in our study of human and mouse outer retina synapses, each dataset consisted of hundreds of synapses and was acquired with a 24-hour turnaround from sample collection to image ready tissue, followed by approximately 8 hours of imaging time per species. RAIN-STORM is thus uniquely suitable for large sample numbers and the acquisition and quantitative analysis of extensive datasets

While improvements in accessibility, speed, and target compatibility were major motivations for the development of RAIN-STORM, several optimized parameters were uncovered that could be useful in other nanoscopic imaging applications. Toward this end, we report quantitative imaging metrics for all 125 tested parameters as a resource for the community. Among these, some unexpected advantages of various conditions were discovered. For example, including a post-fixation step once primary and secondary antibody staining had been performed improved resolution by ∼20nm. We also found that the ratio of BME to MEA could be used to tune fluorophore switching properties, and that BME (140 mM) combined with MEA (40 mM) was most suitable to induce balanced blinking of fluorophores in both imaging channels. Given these results, we anticipate that many of the parameters that we report will be useful to other SMLM approaches.

RAIN-STORM also offers new opportunities for the discovery of novel nanoscopic structures or molecular components. A diverse array of targets could be readily visualized across a range of cell types and subcellular structures, with ∼90% of tested commercial antibodies showing good labeling and resolution. By comparison, most current 3D STORM methods have been developed using a handful of molecular targets (<5), many of which are cytoskeletal (Chamma et al., 2016; Mikhaylova et al., 2015; Mönkemöller et al., 2015; Stahley et al., 2016; van den Dries et al., 2013). RAIN-STORM is thus suitable for biologic discovery of new nanoscopic structures or mapping the distribution or localization of a diverse array of molecular components.

RAIN-STORM may also be useful for the study of molecules and structures beyond those described here. As with other SMLM methods, the relative density of a given protein and the specificity of available antibodies are key factors in determining good targets for RAIN-STORM. As expected, the relative density of an individual target influences the degree of structural filling. Because single molecules are visualized with this approach, the relative labeling density of a given target can report on the location and area of individual substructures, as we demonstrated for ribbon synapses. For imaging the shape and structure of individual cell types, the relative density of a given protein and the availability of high-quality antibodies can make some targets more useful than others. For example, PKCα densely and specifically labels rod bipolar cells and reliably reveals the entirety of the bipolar cellular architecture at the nanoscale, while Iba1, a microglia marker, appears less dense and fills microglia structures incompletely. Notably, RAIN-STORM parameters performed consistently well across molecular targets and species, suggesting RAIN-STORM methods should provide a good baseline for imaging of additional molecular targets and tissue types not tested here.

The many technical advantages of RAIN-STORM enable access to important aspects of glia and neural circuit molecular architecture that were previously difficult to visualize and measure within tissue. For instance, RAIN-STORM provides easy and rapid quantification of individual PSD95 and Ribeye labeled synapses in 3D volumes of neuropil in both mouse and human clinical specimens. Using this approach, we were also able to validate the existence of IP-TNTs and map new molecular features of these structures. This includes the presence of a what appears to be a solid collagen IV core filament that may provide stability to these small structures. Future studies will shed light on how this collagen filament resides within the nanotube and whether it participates in IP-TNT-mediated neurovascular coupling (Alarcon-Martinez et al., 2020). In addition, we documented ∼100-200nm filaments arising from astrocytes in the ganglion cell layer. Various roles for retinal astrocytes have been described, including vascular patterning (O’Sullivan et al., 2017; Tao & Zhang, 2016), but our data hint that they may also physically interact with and perhaps modulate adult retinal neurons through nanoscale structural interactions (Arizono et al., 2020). Given that changes in the numbers and densities of synapses, neurovascular coupling, and in neuron-glia interactions are thought to be central to many neurological and cognitive disorders (Lian and Zheng, 2016; Kempuraj et al., 2018; Verdugo et al., 2019), the potential of rigorously and rapidly measuring such features is an important advance. In addition, because RAIN-STORM is readily compatible with human tissue, this method will also be useful for clinical specimens and comparisons between human disease and animal models. Finally, since RAIN-STORM can be leveraged together with genetic reporters and viral labeling strategies to obtain directed co-imaging of various targets, these methods may offer additional combinatorial advances for overlaying molecular information on cellular ultrastructure. Such investigations of neural circuits and other tissues will help continue to drive discovery of novel nanoscale biological features in native tissue environments in healthy and diseased states.

## Materials and Methods

### Coverslip Preparation

Small batches of #1.5H 22mm round glass coverslips (Neuvitro #GG-25-1.5H) were cleaned via sonication. Coverslips were then submerged in 5M sodium hydroxide (Fisher Scientific #1310-73-2) for 30 minutes and serially washed three times in deionized water. Coverslips were next washed in serial dilutions of ethanol (70%, 90%, 100%) for 30 minutes each and then allowed to dry. Once dry, coverslips were dipped in a solution of 0.5% (m/v) Gelatin A, 7mM Chromium(III) Potassium Sulfate in PBS and again were allowed to dry again before storing up to one month at room temperature.

### Experimental Model and Subject Details

For mouse tissue, retinas were collected from four 6 to 8-week-old C57BL/6J animals. Experiments were carried out in male and female mice in accordance with the recommendations in the Guide for the Care and Use of Laboratory Animals of the NIH under protocols approved by the BCM Institutional Animal Care and Use Committee. Macaque (N = 2), rabbit (N =2), and pig (N = 3) retinal tissue were obtained via the Baylor College of Medicine Center for Comparative Medicine veterinary from unrelated surgical procedures. Human donor eyes (N = 2) were obtained in collaboration with the Lions Eye Bank of Texas at Baylor College of Medicine. Informed consent was acquired from all patients and/or participating family members in accordance with EBAA and FDA regulatory standards. Subjects were 40 and 58 years old and had no documented history of eye disease.

### Tissue preparation

Details for tissue preparation and staining methods are provided for the optimized RAIN-STORM protocol. Tested variations on this protocol are listed in **Supplemental Table 1**. Briefly, tested parameters included both 25°C and 4°C fixations, PFA concentrations of 1%, 2%, and 4%, Glutaraldehyde preparations with PFA (2% PFA, 0.3% GA), or glutaraldehyde by itself (0.3% GA), each prepared in phosphate buffered saline (PBS), with fixation times of 30, 60, or 120 minutes. Quenching conditions included ammonium chloride (10mM, 100mM), glycine (10mM, 100mM) or sodium borohydride (0.1%, 0.5% w/v). For the final selected tissue preparation condition, mouse eyes were enucleated and placed in 2% paraformaldehyde for 1 hour at 4°C, then subsequently rinsed in 100mM glycine solution for 1 hour at 4°C. Samples were then washed in PBS for 30 minutes and stored in PBS. Eye cups were then dissected, removing the cornea and the lens. Samples were then allowed to fully equilibrate in 30% sucrose until the tissue sank (∼45-60 minutes). Tissue was serially washed by hand in Optimal Cutting Temperature (OCT) compound to remove excess sucrose and subsequently placed into molds filled (OCT) compound (Sakura, Torrance, CA). Embedded tissue was then frozen using methyl butane chilled on dry ice. Human eyes were prepared using our final selected fixation condition of 2% PFA at 4°C but were fixed for 4 hours given the increase in tissue thickness. All other conditions were identical to those detailed above. All blocks were then stored at −80°C until ready for use. Tissue was sectioned at 10µm and mounted on prepared coverslips.

### Antibody Staining

For quantification and final images, slides were incubated in the optimized blocking solution (5% normal donkey serum and 0.3% Triton X-100 in PBS) for 1 hour and then with primary antibodies diluted in blocking solution (Table 1) for a minimum of 12 hours at 4°C. Other tested conditions include variations in concentrations of Triton-X100 (0.1%, 0.3%, 0.5%, 1.0%, 2.0%), Saponin (0.1%, 0.3%, 0.5%, 1.0%, 2.0%), and Normal Donkey Serum (1%, 3%, 5%, 10%, 15%). Samples were washed with PBS three times for 20 minutes and then incubated with commercial dye-conjugated secondary antibodies diluted in blocking solution (AF647-conjugates from Jackson ImmunoResearch Laboratories, West Grove, PA, and CF568-conjugates from Biotium, Fremont, CA) for 1 hour at room temperature. Slides were then washed with PBS three times for 20 minutes prior to applying a 4% PFA solution as a postfix for 30 minutes. Slides were washed with PBS three times for 20 minutes each, 100mM NH_4_Cl was applied for 30min, and slides were again washed with PBS three times for 20 minutes each. Samples were stored in PBS until imaging.

### Imaging Buffer

Imaging buffer for use in STORM was prepared using stock solutions of one of three oxygen scavenging enzymes: pyranose oxidase (0.5 U/µL, Millipore Sigma #P4234-250UN), glucose oxidase (Millipore Sigma #G2133-50KU), or protocatechurate 3,4-dioxygenase (PCD, Millipore Sigma #P8279-25UN). Each oxygen-scavenging enzyme was used to name the associated buffer formulation. Tested imaging buffers also contained bovine-derived catalase (100 U/µL, Millipore Sigma #C1345-10G), cysteamine hydrochloride (MEA, 1 M, Chem-Impex International #02839), and 2-mercaptoethanol (BME, Millipore Sigma #M6250-250ML). When PCD was used, its substrate protocatechuic acid (PCA, 3,4-dehydroxybenzoic acid, Millipore Sigma #37580-25G-F) was also included. For a given buffer formulation, one of the components was varied by concentration, keeping all others constant. Once this was performed for each buffer-enzyme combination, the final selected buffer was tested with either cyclooctatetraene (COT, Millipore Sigma #138924-1G) or trolox (Millipore Sigma #238813-1G) in varied concentrations (1mM, 2mM, or 5mM). Stock solution aliquots were kept frozen at −80°C until just prior to use. Aliquots were thawed at room temperature and added to a freshly prepared solution of 30% (m/v) glucose in PBS. Once prepared, imaging buffer was allowed to equilibrate for 20 minutes and then used.

### Imaging and Image Processing

Image acquisition was performed on a Bruker Vutara 352 (Bruker, Billerica, MA) using a water objective (UPLSAPO60XW). Stained samples were mounted in a collared well (Thermofisher Scientific #A7816), and 1mL of imaging buffer was added on top in an open well configuration. All images were acquired at 200nm axial steps using a framerate of 67 Hz for AF647 and 40Hz for CF568 across the full 10µm thickness of tissue using three sequential cycles of 250 frame-captures for each channel, yielding a total of 38250 frames per probe per sample. Between each cycle a timed pause was included to allow for the imaging buffer to re-equilibrate. 3D nanoscopic imaging volumes were thus 40µm by 40µm by 10µm (16,000 µm^3^). For the 640nm, 561nm and 405nm excitation lasers, laser powers of ∼98-110mW (6125-6875 W/cm^2^), ∼90-100mW (5625-6250 W/cm^2^), and ∼2-6mW (125-375 W/cm^2^), were measured at the sample using a 40µm by 40µm field of view. Data were then clustered to determine associated and contiguous structures within the image using the Ordering Points to Identify the Clustering Structure (OPTICS) algorithm. To analyze images, a general particle distance of 0.16 µm and a particle count of 25 was used for all channels on all images. The particle distance refers to the maximum allowed distance that a particle can be from another particle in order to be included in a given cluster while the particle count reflects the minimum required number of particles required to form a cluster. Any non-clustered localizations were removed from the image. Once filtering and clustering were complete, an FRC analysis was performed to determine the global aggregate resolution of the sample in the XY and XZ dimensions for each target imaged using three repeat measurements for curve smoothing.

### Data availability

All data are available on reasonable request from the authors.

---

**Video 1: RAIN-STORM-imaged horizontal cell shows striking neuronal arbor detail.** Tissue was prepared for RAIN-STORM, and Calbindin-labeled horizontal cells were imaged across 10µm. The image is colored depth by depth, with colors indicating the axial position throughout the stack (blue 0µm to yellow 10µm). Fine neurite structural detail is observed, and individual synapses are clearly visible.

**Video 2: RAIN-STORM-imaged blood vessel shows an interpericyte tunneling nanotube bridging two vessels.** Tissue was prepared for RAIN-STORM, and collagen IV-labeled vessels were imaged across 10µm. The image is colored depth by depth, with colors indicating the axial position throughout the stack (blue 0µm to yellow 10µm). The structure and morphology of hollow vessels are well preserved, and cross-vessel nanotube connections are apparent.

**Video 3: RAIN-STORM-imaged astrocytes demonstrate fine filamentous structures and a mesh of interacting fibers.** Tissue was prepared for RAIN-STORM, and GFAP labeled astrocytes were imaged across 10µm. The image is colored depth by depth, with colors indicating the axial position throughout the stack (blue 0µm to yellow 10µm). The structure and morphology of astrocyte filamentous fibers was observed, which appear to form a highly branched network.

**Video 4: RAIN-STORM imaging of mouse bipolar neurons and ribbon synapses reveals pre- and postsynaptic neural interactions**. Tissue was prepared for RAIN-STORM, and mouse bipolar neurons in the outer plexiform layer were labeled with PKCα (magenta) and pre-synaptic ribbons were labeled with RIBEYE (cyan). The image is colored depth by depth, with colors indicating the axial position throughout the stack (blue 0µm to yellow 10µm). Individual neurite tips can clearly be observed invaginating individual presynaptic ribbons.

**Video 5: RAIN-STORM imaging of human bipolar neurons and ribbon synapses shows bipolar cell interactions with ribbon synapses.** Human retina was prepared for RAIN-STORM, and human bipolar neurons in the outer plexiform layer were labeled with PKCα (magenta) and pre-synaptic ribbons were labeled with RIBEYE (cyan). The image is colored depth by depth, with colors indicating the axial position throughout the stack (blue 0µm to yellow 10µm). As in mouse sample, individual human bipolar neurite tips interact with and invaginating individual presynaptic ribbons. While fewer synapses were observed in the human samples relative to the mouse, it is likely that this decrease is due to the inherent delay in human post-mortem sample collection rather than a biologically relevant difference.

**Figure 1 – figure supplement 1:**
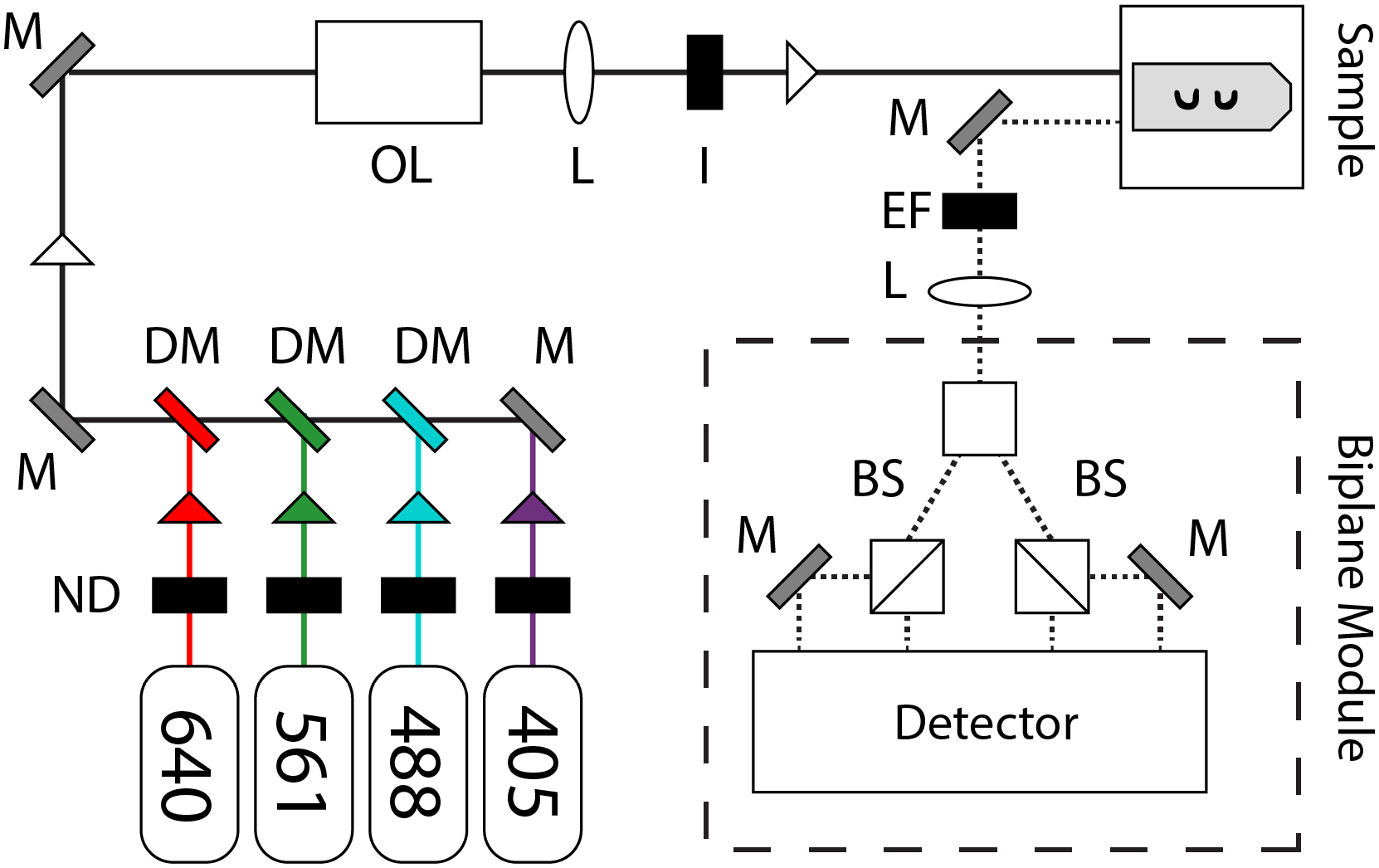
Optical Diagram of Vutara. Layout and optical design of the Bruker Vutara SRX352, which allows 3D STORM imaging and PSF localization via a biplane module in place of a cylindrical lens. **M**: Mirror, **DM**: Dichroic mirror, **ND**: Neutral density filter, **L**: lens, **BS**: Beam splitter, **I**: iris/aperture, **OL**: Objective lens, **EF**: Emission filter.

**Figure 1 – figure supplement 2:**
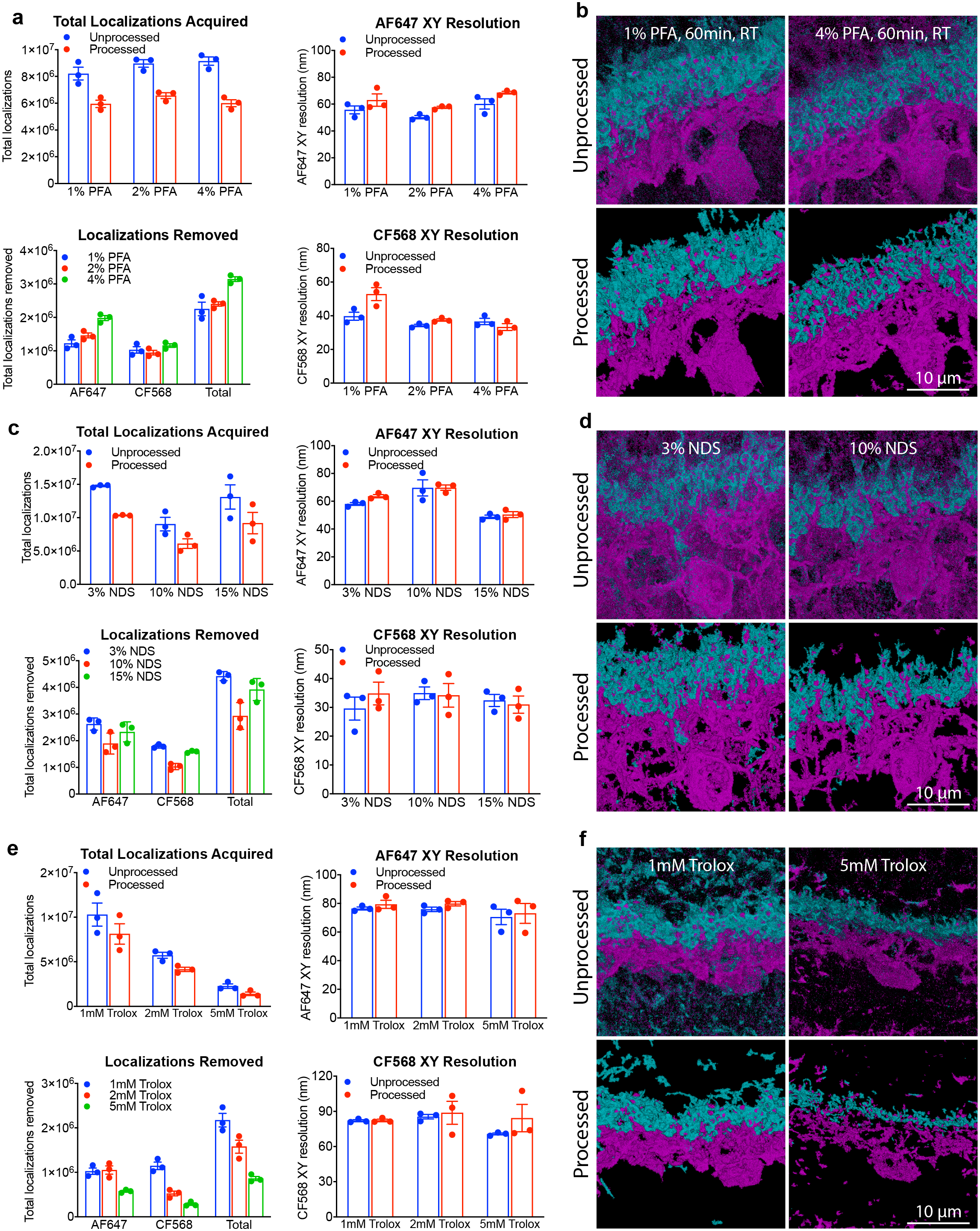
Modifying sample conditions improves visual quality and image metrics. **a-b**, Representative primary data metrics (**a**) and corresponding unprocessed and processed images (**b**) are shown for variations in primary fixation concentrations. A small increase in total localizations acquired was observed with increasing concentrations of PFA, though the gain in localizations was offset by background localizations at these concentrations. Based on these parameters, 2% PFA was selected for optimized imaging. **c-d**, Representative primary data metrics (**c**) and corresponding unprocessed and processed images (**d**) are shown for variations in blocking buffer serum concentrations. In general, the best image metrics were obtained for mid-level serum concentrations (e.g. 3-5% NDS), while the lower (e.g. 1% NDS) and higher (e.g. >10%) serum concentrations resulted in either increased filtered localizations or decreased resolution overall and poorer image quality. **e-f**, Representative primary data metrics (**e**) and corresponding unprocessed and processed images (**f**) are shown for variations in imaging buffer formulations using Trolox. Increasing the Trolox concentration reduced the total amount of data collected resulting in poorer image quality relative to low Trolox concentrations (e.g. 1mM). Images are representative of those acquired from N=3 animals. Calbindin, magenta; PSD95; cyan. Scale bars = 10 µm. Data are represented as the mean ± the s.e.m. Both unprocessed (blue) and processed (red, OPTICS algorithm) data are shown for each dataset.

**Figure 1 – figure supplement 3:**
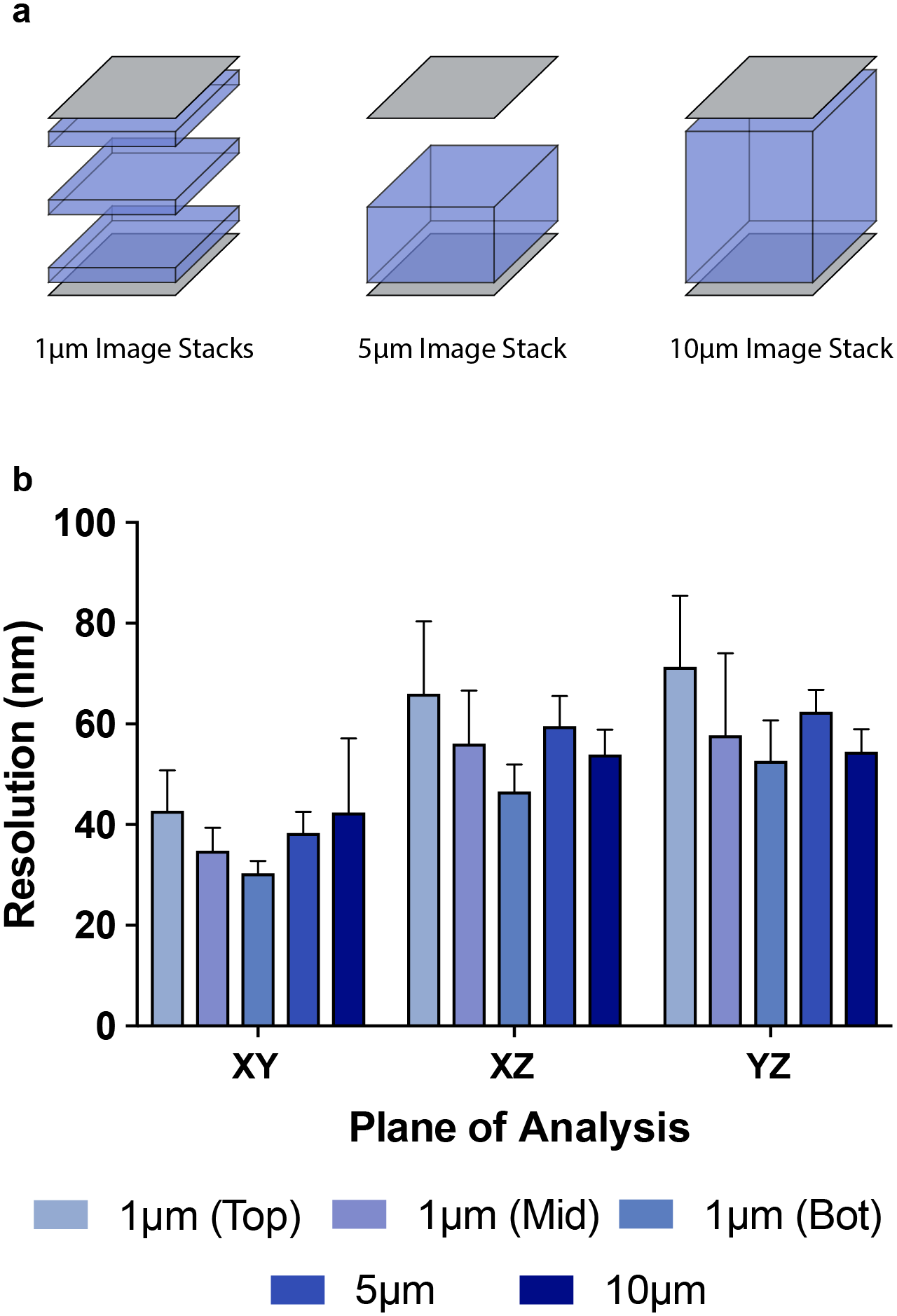
RAIN-STORM resolution as a function of sample depth. **a**, Schematic of three image planes used to calculate sample resolution. Resolution was compared in: 1) single-micron slices at the bottom of the image (nearest the objective), the middle of the image stack, or at the top of the image; 2) 5µm slices from the bottom of the image to the midpoint; and 3) the entirety of the 10µm stack. **b**, Sample resolution quantification as a function of the three calculation methods schematized in (**a**). Planar resolution (XY dimension) varies from 30.0 ± 1.6 nm to 42.5 ± 4.8 nm while resolution in the axial dimensions (XZ and YZ) ranges from 46.3 ± 3.3 nm nearest the objective to 65.7 ± 8.5 nm farthest from the objective. N = 3. Data are represented as the mean ± the s.e.m.

**Figure 2 – figure supplement 1:**
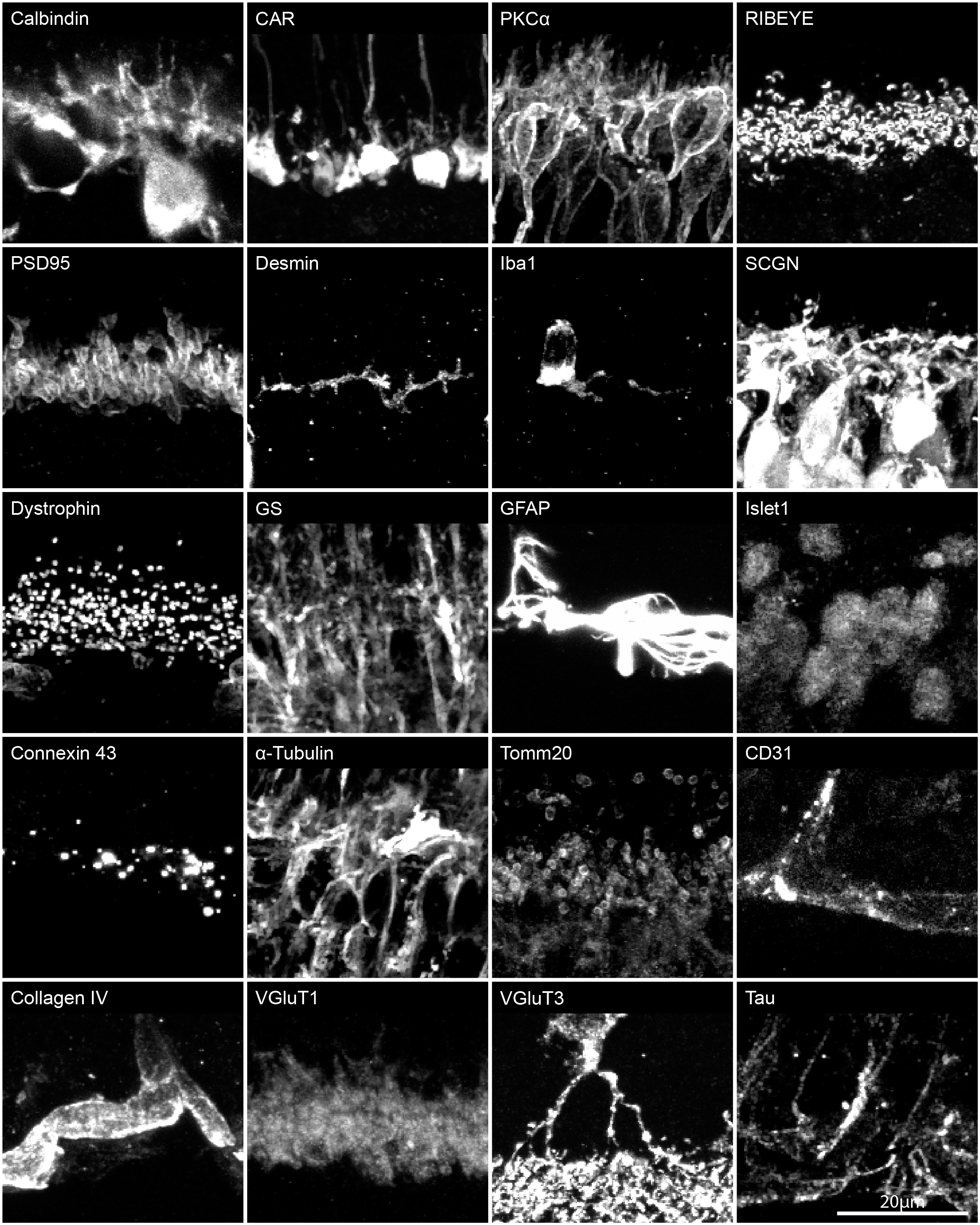
Confocal imaging verifies antibody specificity in RAIN-STORM imaging. Representative confocal images of antibodies used in this study. Cellular structure and labeling patterns from confocal imaging were used as a baseline with which to compare the effectiveness of RAIN-STORM imaging for various targets. Scale bars = 20 µm.

**Figure 2 – figure supplement 2:**
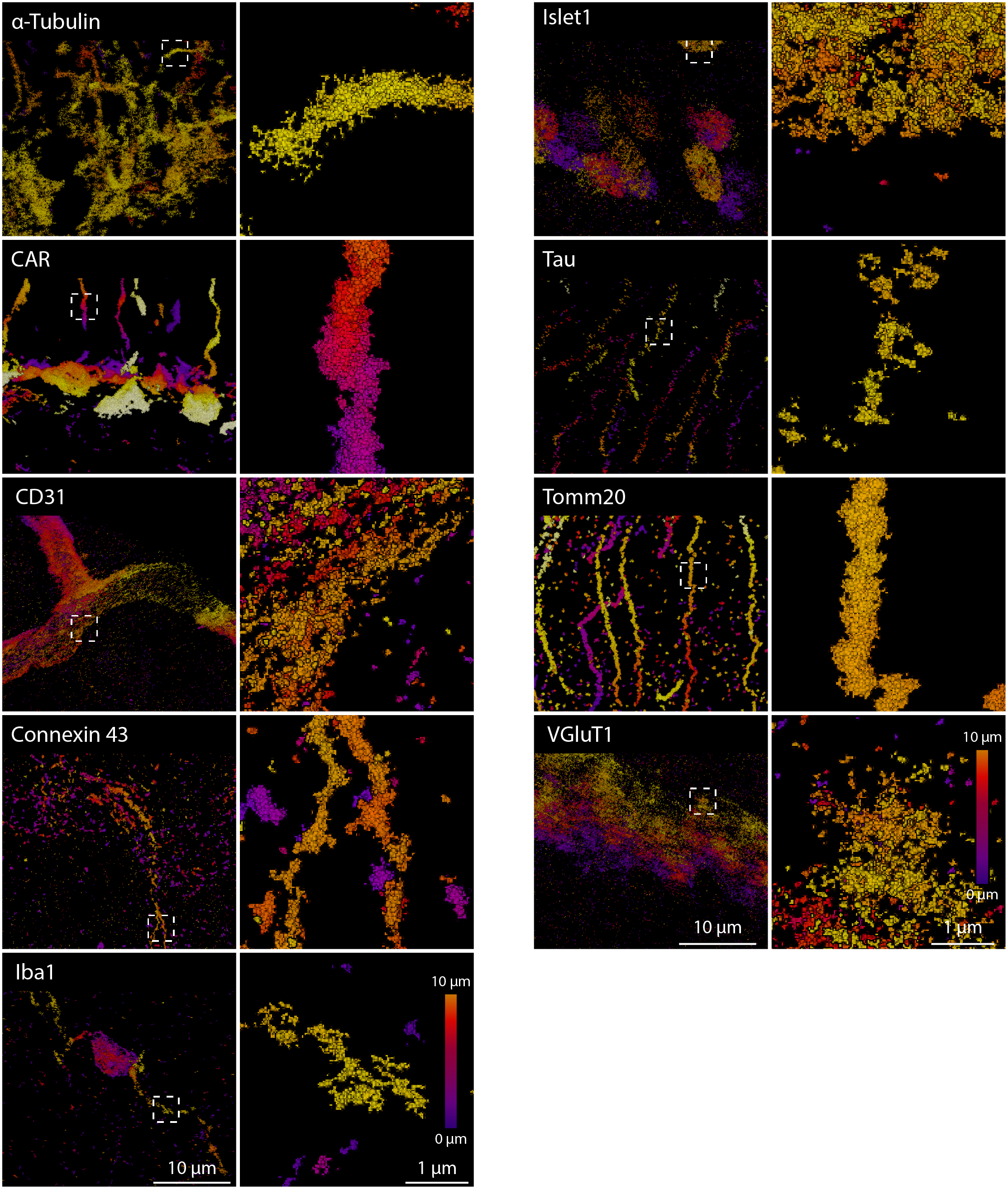
RAIN-STORM imaging can be applied to diverse array of molecular targets. RAIN-STORM imaging of α-Tubulin, a marker for cytoskeletal tubulin in the murine retina, cone arrestin, a marker for cone photoreceptors, CD31, a marker for select blood vessel structures, Connexin 43, a gap junction protein, Iba1, a marker for microglia, Islet1, which stains a variety of neuron cell bodies, Tau, a microtubule-associated protein, Tomm20, a mitochondrial marker, and VGluT1, a glutamate transporter marker for specific subsets of cells. These stains demonstrate a wide variety of protein types and targets that are amenable to imaging with RAIN-STORM. N = 3 animals. Scale bars = 10 and 1µm.

**Supplemental Table 1:**
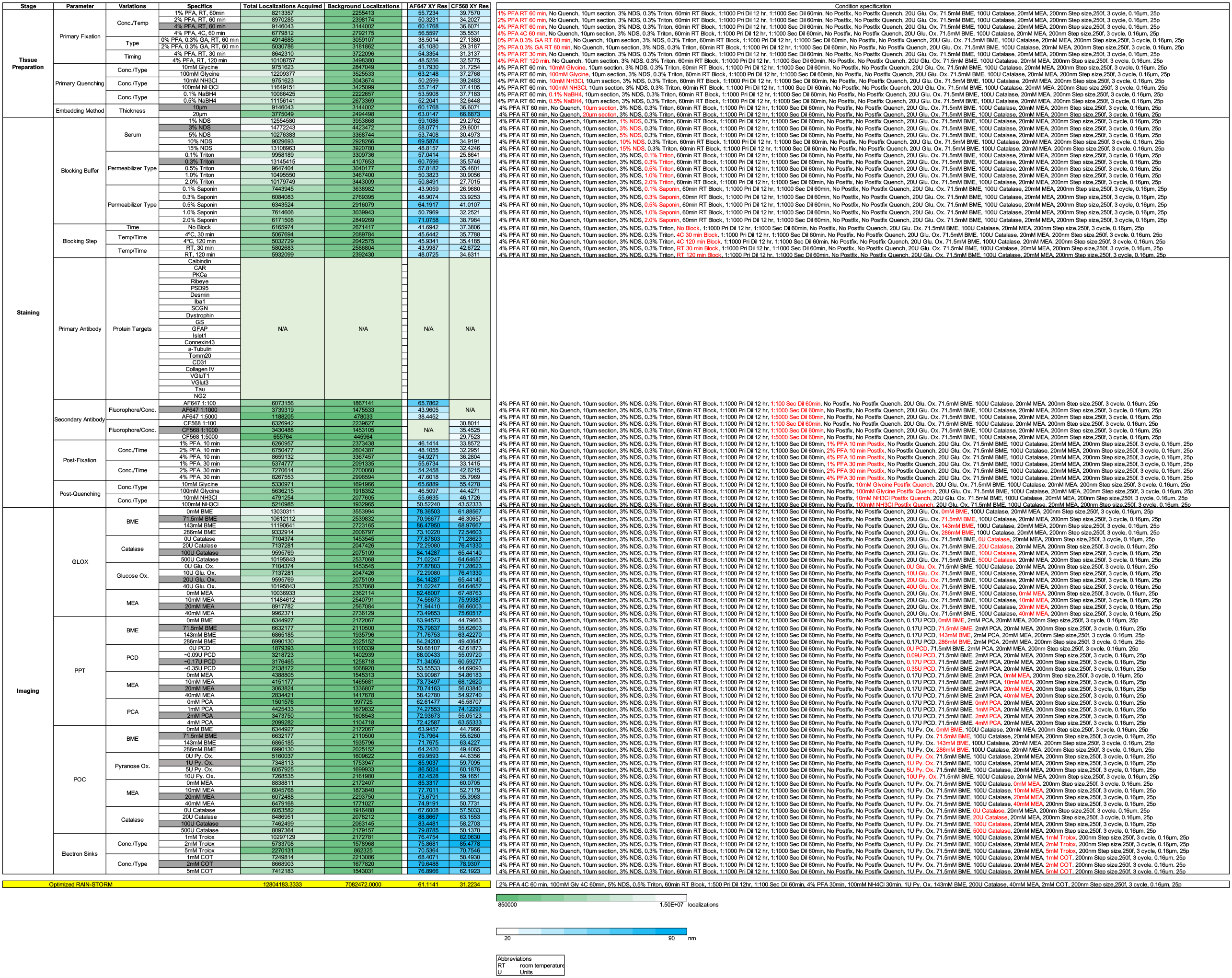
Summary of condition variations tested for RAIN-STORM.

**Supplemental Table 2:**
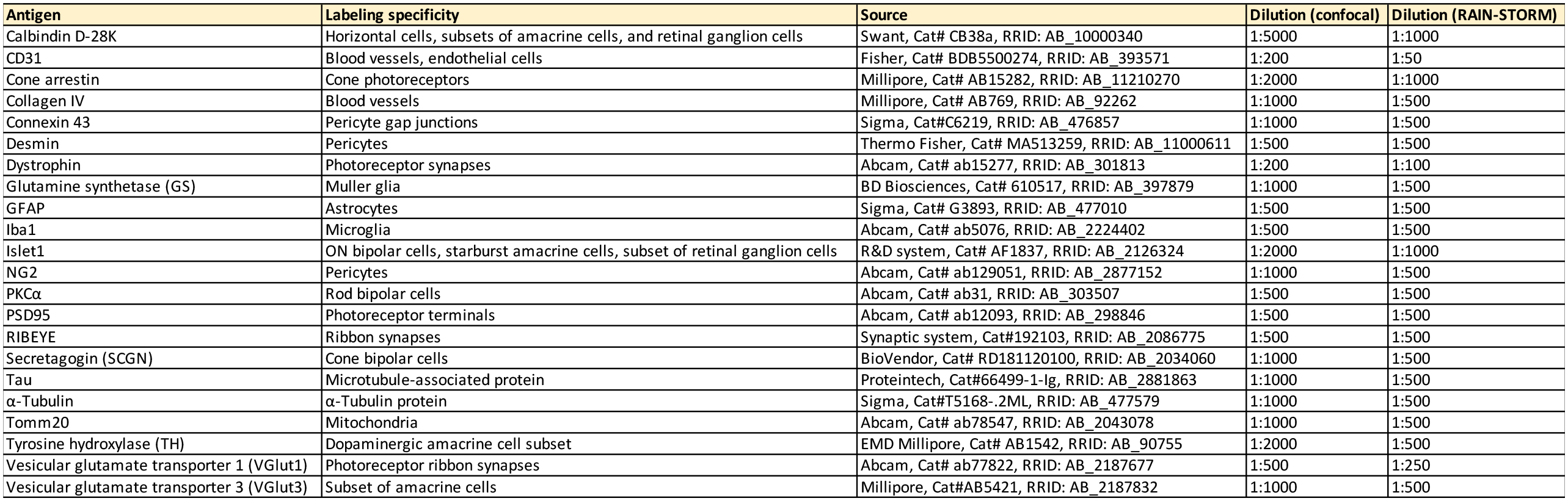
Primary antibodies used.

